# Towards sequential conjugation-assisted laboratory evolution (SCALE) of Cas nucleases

**DOI:** 10.1101/2024.03.20.585880

**Authors:** Thomas Swartjes, Kristel Doets, Raymond HJ Staals, John van der Oost

## Abstract

DNA endonucleases such as the CRISPR-associated enzymes Cas9 and Cas12a constitute powerful molecular tools with a wide range of applications. To facilitate optimization of these enzymes, we investigated a fully *in vivo* selection system for laboratory evolution based on plasmid conjugation. In the proposed method, DNA cleavage promotes plasmid transfer to other cells, allowing cleaving variants to spread through the bacterial culture via consecutive conjugations. We have investigated whether the conjugation system of the F plasmid of *E. coli* is suitable for laboratory evolution of Cas nucleases. Through a series of conjugation experiments, we found (i) that the F plasmid can spread through bacterial cultures, (ii) that the FinOP conjugation inhibition system is a potential nuclease target as it can prevent transfer of non-cleaving variants (counterselection), and (iii) that a trimmed version of the F plasmid allows for consecutive conjugations. Overall, we were able to show that in many regards, conjugation can indeed be used as means of *in vivo* laboratory evolution. We discuss the potential of the here presented concept for a yet to be developed laboratory evolution platform.

## Introduction

### Genome editing

Genome editing holds great promise for various applications. It can be used to improve crop plants (1–3), to cure genetic diseases (4), to empower biotechnological production of compounds (5), and to accelerate fundamental genetic research (6, 7). Genome editing can be achieved by introducing a double stranded break at a specific location in the DNA sequence where editing is desired. Endogenous cellular DNA repair is then responsible for creating the actual edit in the DNA, generally through either non-homologous end joining (NHEJ) or homology-directed repair (HDR) (8). Resolving DSBs through NHEJ results in short insertions or deletions (indels) (9), while HDR uses a repair template of which the flanks are homologous to the ends surrounding the DSB, allowing for a double cross-over to repair the DSB (8).

For genome editing, zinc-finger nucleases (ZFNs) (10, 11) and transcription activator-like effector nucleases (TALENs) (12–14) have been used to introduce DSBs at the desired DNA sequences. The challenge with ZFNs and TALENs is that if a new DNA region is to be targeted, multiple amino acids of these proteins must be adapted to bind and cleave the new DNA sequence. This limitation of ZFNs and TALENs is largely overcome by new molecular tools based on CRISPR-Cas (clustered regularly interspaced short palindromic repeats, and CRISPR-associated genes/proteins). The specificity of Cas proteins Cas9 and Cas12a, can easily be adapted by introducing an RNA guide that is complementary to the desired DNA target.

### CRISPR-Cas

CRISPR-Cas has evolved as an adaptive immune system of prokaryotes to protect themselves against invading DNA and RNA from phages and plasmids (15–17). It provides immunity in three stages: adaptation, expression, and interference (18). During adaption, so called protospacer sequences in the DNA or RNA from the invading are recognized and incorporated in a chromosomal locus called the CRISPR array, where these sequences – now called spacers - are stored between repeat regions (19). In the expression stage, the CRISPR array is transcribed as a precursor CRISPR RNA and subsequently processed into guide RNAs (gRNAs), each carrying a single invader-derived sequence flanked by parts of the repeat. The gRNAs form a complex with a single or multiple Cas protein(s). The resulting ribonucleic protein (RNP) complex of DNA-targeting Cas systems scans the DNA for protospacer adjacent motifs (PAMs) and probes whether the adjacent sequence matches to the gRNA. Sequences with both a PAM (20) and complementarity to the gRNA (15, 16) are then targeted for interference, generally through DSB formation (21, 22).

Cas effector proteins like Cas9 and Cas12a are DNA endonucleases and can be used to create DSBs at specific DNA sequences. For each target, a synthetic gRNA should be designed to control where the nuclease cleaves. Although the targeting range is still limited by the requirement of a PAM, Cas nucleases are more easily adapted to target new sequences compared to ZFNs and TALENs and thus provide a powerful tool for genome editing.

Cas nucleases have also been engineered for other approaches of genome editing, which do not rely on DSB-induced DNA repair. For instance, through mutations in the nuclease domain(s) of Cas nucleases, variants have been created that still bind DNA complementary to the gRNA, but that no longer cleave (catalytically inactive variants) or only cleave one DNA strand (nickase variants) (23). Some of these variants have since been turned into base-editors by fusing them with deaminase domains from other proteins to catalyze base conversions in (or close to) the targeted DNA or RNA (24–27). In addition, a Cas9 nickase variant has been combined with reverse transcriptase and an extended gRNA to yield a prime editor, that is used either to make small alterations (insertions, deletions, or replacements) around the nick site, using the extended gRNA as a template (28), or rather to introduce larger changes (29).

Together, these tools provide great options for genome editing. However, there are challenges still to address. For instance, the fact that PAM-requirements of Cas nucleases can limit some applications has stimulated the generation of variants with adjusted PAM specificity (30–33). Another issue that needs attention is the development, both in mice and humans, of immunity against Cas nuclease proteins (34–36), hampering the use of Cas nucleases for therapeutic genome editing. In addition, when a Cas nuclease and guide are to be delivered in a patient’s body using adeno-associated virus (AAV) vectors, the large size of Cas9 and Cas12a can be problematic as AAVs have a limit to how much cargo DNA they can deliver (37). This especially complicates the delivery of base-editors, prime-editors, or other fusion proteins as these require an even larger coding sequence. Many of these and other challenges are very specific to the intended application. Therefore, it would be ideal to have a system to adapt Cas proteins rapidly and easily for their intended use case.

### Laboratory evolution

Customization of these proteins would benefit from a suitable platform for laboratory evolution. In laboratory evolution, a gene of interest (GOI) undergoes multiple rounds of mutagenesis, expression of the encoded protein (protein of interest, POI), selection based on POI activity, and amplification of the selected GOI variants. Each of these steps can take place either *in vitro* or *in vivo*. A laboratory evolution platform that works completely within living cells with continuous *in vivo* mutagenesis would allow many evolutionary cycles while requiring minimal manual interference. Continuous *in vivo* mutagenesis could, for instance, be achieved using a mutagenesis plasmid (38, 39). This mutagenesis plasmid can substantially increase the mutation rate *in vivo* by preventing export of mutagenic nucleobases, by cytidine deamination, and by impairing DNA proofreading. Another advantage of *in vivo* laboratory evolution is that *in vivo* selected nucleases are likely to function well in physiological conditions.

An established approach for *in vivo* laboratory evolution uses bacteria that express the GOI from a plasmid in such a way that the desired activity leads to improved bacterial growth (40, 41). For example, ornithine cyclodeaminase was evolved towards increased production of L-proline in cells with an artificial growth defect, where the additional L-proline rescued cell growth (42). In combination with continuous *in vivo* mutagenesis, growth-coupled selection may allow multiple evolutionary cycles without manual interference, until the activity of the POI is no longer limiting. When selecting for DNA cleavage, this is problematic because cleavage of DNA in one cell would also affect all future offspring. As a result, DNA cleavage would promote growth in all subsequent generations of daughter cells, regardless of whether the GOI still displays the desired activity. This would effectively limit this approach to just a single evolutionary cycle when DNA nucleases are involved.

One way in which this hurdle has been addressed, is by modifying the GOI beforehand to encode a catalytically inactive protein variant (23), which binds DNA but does not cleave it (33). This variant is then evolved while using DNA binding as a representation for the desired DNA cleavage or modification. The variant would for instance be challenged to bind the promoter of a gene that inhibits growth. Stable binding can inhibit gene expression (43, 44), indirectly promoting cell growth. After laboratory evolution, the mutations that made the GOI catalytically inactive would be reverted, assuming that the GOI is now better suited to cleave or modify the target DNA. However, the evolved protein may have accumulated mutations that disturb the desired activity (DNA cleavage) since only variants with enhanced DNA binding were selected for (“you get what you screen for”) (45). It would therefore be more effective to select for the activity that is required in the end. For DNA cleaving enzymes like Cas9 and Cas12a, this would require a way to refresh the DNA substrate after each round of selection.

### Horizontal gene transfer in laboratory evolution

This limitation could be addressed by using horizontal gene transfer (HGT) of either the DNA substrate or the GOI itself. The latter has been successfully shown in phage-assisted continuous evolution (PACE), a laboratory evolution platform where the GOI is expressed from a modified M13 bacteriophage (38). In short, the desired activity of the POI induces expression of M13 *gene III*, which encodes a phage coat protein that is required for the production of new intact M13 phages. PACE has been used to evolve a Cas9 variant that recognizes a broader range of PAMs (33). Although PACE is an elegant and powerful platform, it can be difficult and laborious to set up and optimize (46).

Besides viruses, as used in PACE, HGT can also be mediated by (i) transformation, where cells take up DNA from the environment, or (ii) conjugation, where plasmid DNA is transferred from one cell to another. The best understood plasmid that causes its own conjugative transfer is the F plasmid, which expresses most genes involved in conjugation from a 33 kilobases (kb) long sequence called the transfer region (tra- region) (47) (Figure 1A). To enable conjugation, cells with the F plasmid (donors) construct pili on their surface. These pili extend and bind to cells that do not yet have the F plasmid (recipients) (48). The pili then retract, pulling the recipient cells to the donor cell (49), allowing the formation of a stable mating pair (50). Within the donor, several F-encoded proteins together form a complex called the relaxosome, which then binds (51) and nicks (52) the origin of transfer (*oriT)*, a DNA sequence in the F plasmid (Figure 1A). The two DNA strands of the F plasmid are then unwound, after which the nicked strand is transferred to the recipient cell. As a result, the first region of the F plasmid to enter the recipient cell is directly upstream of the tra-region and is called the leader region (the tra-region itself transfers last). After transfer, the nicked strand is circularized and for both strands, the complementary strand is generated (53). Over time, the recipient cell has been transitioned into a donor cell, able to transfer the F plasmid to other cells (54). The leader likely plays a role in the timely circularization of the plasmid, production of the complementary strand, gene expression, and other processes required for cells to transition from recipient to donor (55, 56).

**Figure 1.**
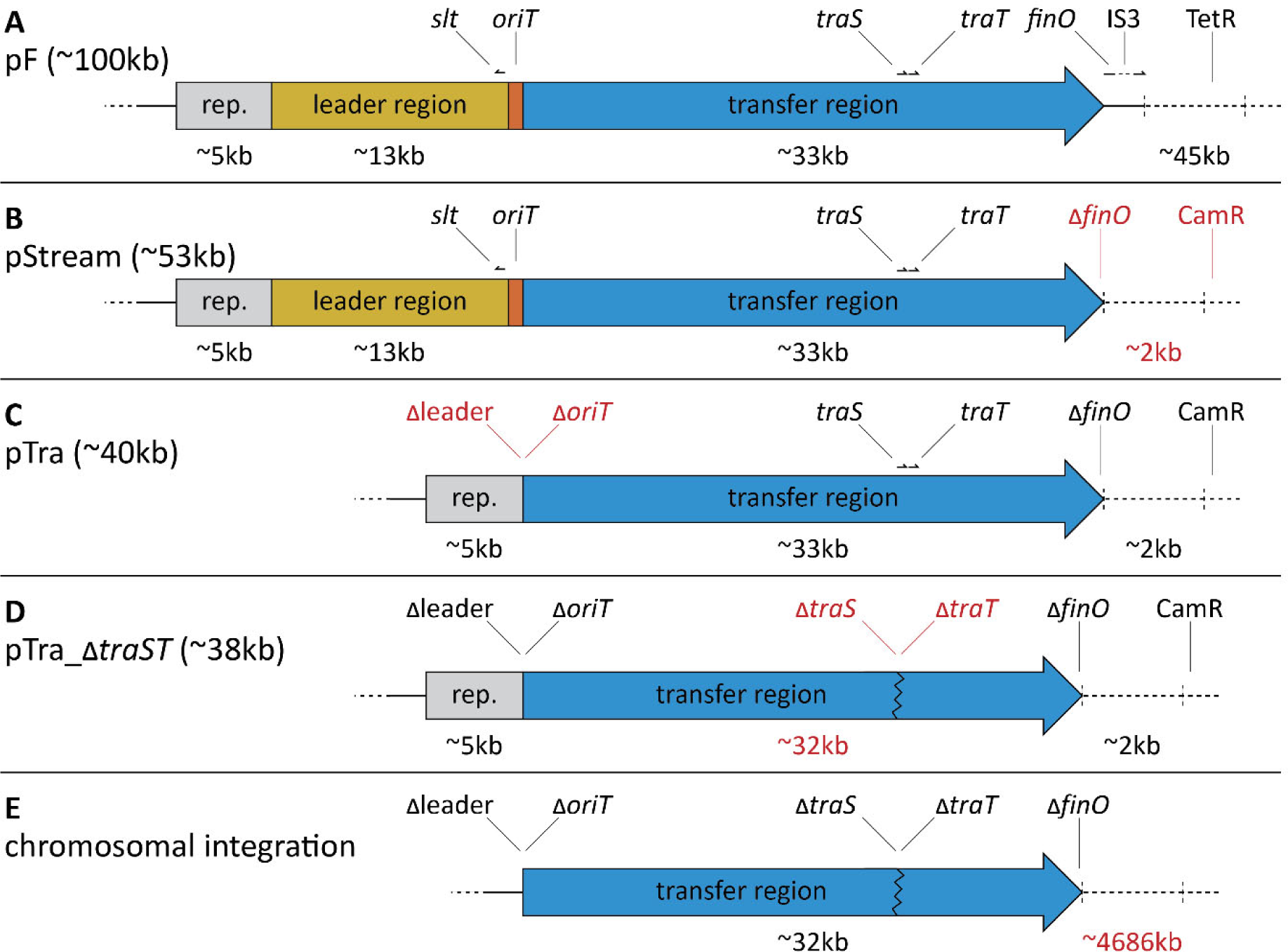
Organization of the F plasmid and derived constructs. Schematic representation of the relevant regions and genes of the F plasmid (pF), and construct based thereon. Elements are shown roughly to scale, except for *oriT* (290bp), which is overrepresented in size for visibility. Changes compared to the above construct are shown in red for emphasis. The plasmids in panels A-D are also included in Supplementary Table 1. **(A)** pF, the F plasmid. **(B)** pStream, a streamlined version of the F plasmid. **(C)** pTra, a plasmid intended to facilitate transfer of a secondary plasmid with *oriT*. **(D)** pTra_Δ*traST*, pTra without *traS* and *traT*. **(E)** The planned integration of the tra- region from pTra_Δ*traST* into the chromosome of *E. coli* DH10B.

In donor cells, the F-encoded protein TraJ induces expression of many genes involved in conjugation (57, 58). TraJ translation is inhibited by antisense RNA FinP and protein FinO (59), both expressed from the F plasmid. The FinO protein stabilizes the FinP RNA (59, 60) and allows it to bind the TraJ transcript (61), inhibiting conjugation. In this manner, the FinOP inhibition system regulates conjugation from within the donor cells. However, the F plasmid also regulates conjugation from would-be recipient cells. If a potential recipient cell already contains the F plasmid, conjugation is prevented. This prevention of conjugation is mediated by F-encoded proteins TraS and TraT, that prevent mating pair formation (62) and DNA transfer (63), respectively. In this manner, the F plasmid prevents unnecessary conjugations, reducing the burden for host bacteria (64, 65).

Overall, the F plasmid constitutes a well-understood platform for HGT, which could potentially be applied to laboratory evolution. Here, we describe a novel HGT-based method for laboratory evolution: sequential conjugation-assisted laboratory evolution (SCALE). We share our efforts to establish this platform based on the conjugative system of the F plasmid, and show that, in many ways, the envisioned steps are feasible. However, some final steps remain to be completed to fully realize SCALE.

## Results

### Design of sequential conjugation-based laboratory evolution

We investigated whether conjugation by the F plasmid could be used to establish a laboratory evolution platform for Cas nucleases. In the envisioned platform, SCALE, a variant of the POI that displays the DNA cleavage, promotes conjugation of the plasmid with the GOI to other cells. These new cells have an unaffected copy of the target DNA, allowing another round of evolution. In this way, only the desired variants can spread through the bacterial population by means of conjugation.

Because conjugation by the F plasmid involves many genes, large constructs are likely required in order to test different parts. To address this, we selected *Escherichia coli* strain DH10B (66) as a suitable strain for the conjugations, because it is well-adapted for maintaining large DNA constructs (67). In the proposed design, DH10B cells would contain all the genes required for F plasmid conjugation, most of which are encoded on the transfer region (tra-region). Eventually, it would be ideal to express these genes from the bacterial chromosome, to minimize the number and size of the plasmids that are needed. By default, conjugation would be prevented due to the presence of a plasmid containing the *finO* gene, which – together with *finP* – inhibits conjugation. The GOI would be expressed from a separate plasmid that contains the *oriT* DNA sequence from the F plasmid. The *oriT* would have allowed this GOI-plasmid to be transferred to other cells if *finO* had not been present.

To start laboratory evolution, the GOI-plasmid is transformed to some of the host cells that already contain the conjugative genes and the *finO*-plasmid (Figure 2, step 1). If the protein encoded by the GOI does not cleave the *finO*-plasmid (Figure 2, step 2a), conjugation remains repressed. The GOI-plasmid would thus be ‘stuck’ within the cell, not able to transfer to other cells. However, if the GOI does cleave the *finO*- plasmid (Figure 2, step 2b), FinO protein production is halted, and FinO proteins will be lost over time. This relieves inhibition, allowing expression of the genes from the tra-region (Figure 2, step 3), causing conjugation of the GOI-plasmid to recipient cells (Figure 2, step 4). The recipient cell now contains both the *finO*-plasmid and the GOI-plasmid (Figure 2, step 5). The GOI is thus challenged again to interfere with the *finO*-plasmid, starting a new round of selection.

**Figure 2.**
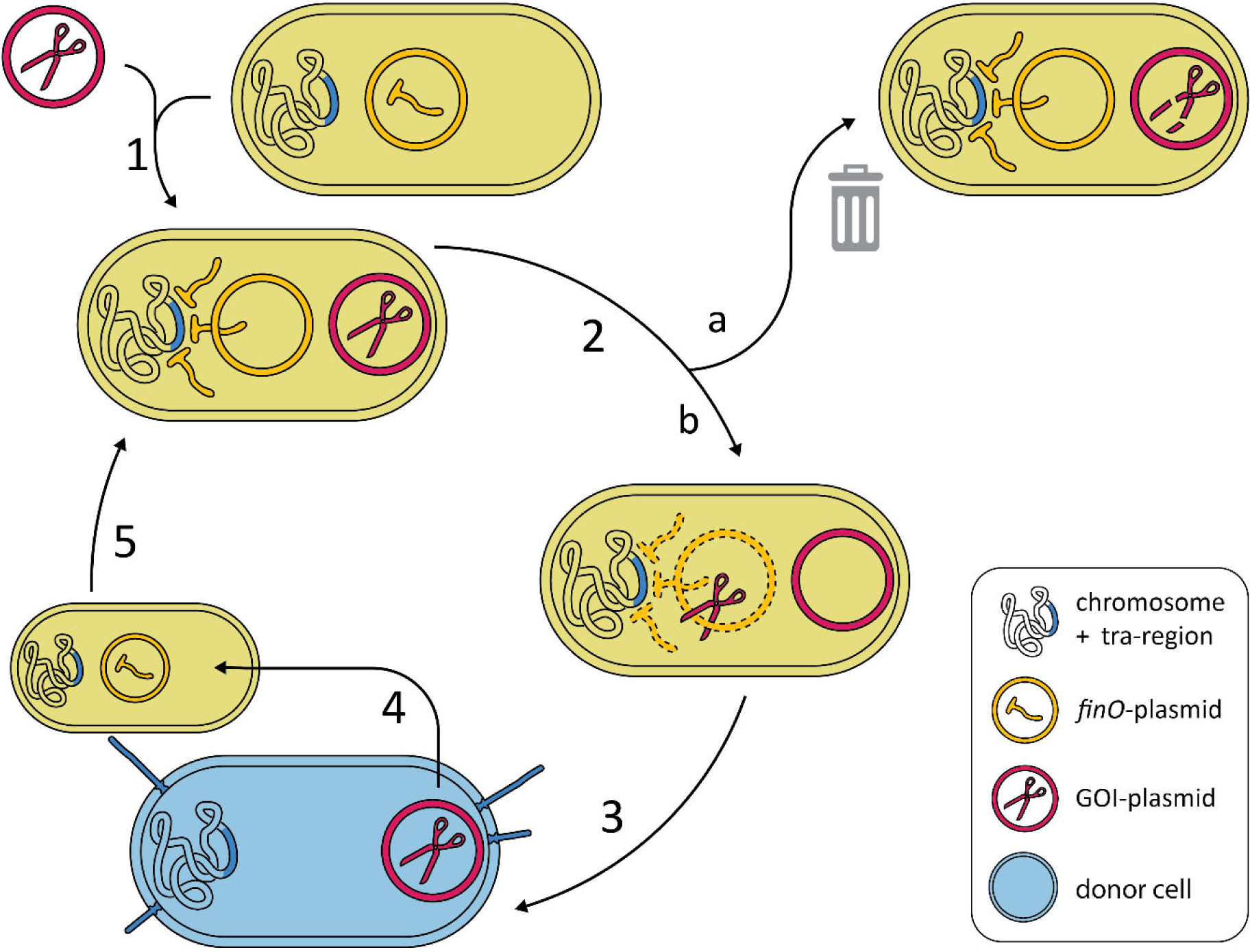
Schematic representation of SCALE. Each cell is depicted with a bacterial chromosome on the left side with, in blue, the genes needed for facilitating conjugation (tra-region). Plasmids are shown as colored rings, either orange (*finO* plasmid) or red (GOI plasmid). The FinO protein is represented by orange T-shapes, while the protein encoded by the GOI is shown as a red pair of scissors. Dashed lines are used to indicate plasmids or proteins that are being lost. The blue cell at the bottom is depicted with pili to signify that it can act as a donor for a subsequent round of conjugation. The envisioned approach is divided in 5 steps. **1:** transformation of host cell with (variants of) the GOI plasmid. **2:** Selection based on the activity (plasmid cleavage) of the protein encoded by the GOI. **2a:** Protein variants without the desired activity are unable to interfere with repression of conjugation. These cells will therefore not contribute to propagation of their GOI variant **2b:** Variant proteins with the desired activity cleave the *finO*-plasmid. **3:** FinO protein is lost over time, relieving repression of conjugation. **4:** The GOI-plasmid transfers to other bacterial cells. **5:** Recipient cells now contain both plasmids and enter a new round of evolution.

In practice, this approach starts with two types of cells: recipients (lacking a GOI-plasmid) and potential donors (with a GOI-plasmid). Plasmids with a functional GOI spread through the culture (by successful conjugation to recipient cells), eventually saturating the culture with cells containing a variant of the GOI- plasmid. At this point, the culture should be diluted with fresh recipient cells, allowing additional rounds of evolution. Finally, after many rounds of enrichment, the GOI-plasmid should be sequenced. Enriched variants can then be tested for the desired activity in a separate assay. When combined with *in vivo* continuous mutagenesis, we hope that the described selection approach enables enrichment of DNA nuclease variants with beneficial mutations, toward their intended applications.

### Conjugation by the F plasmid

To test conjugation by the F plasmid, we generally conducted liquid mating experiments with equal concentrations of donors and recipients. Both donors and recipients were grown in rich medium for two hours, upon which the cultures were diluted to the same optical density (OD_600_). Equal volumes of donors and recipients were then mixed, and subsequently incubated for one hour to allow mating (68). The mating reactions were stopped by vigorous shaking. To confirm that *E. coli* DH10B can act both as donor and as recipient for F plasmid conjugation, we used donor DH10B cells containing the F plasmid, called plasmid pF from here on (Supplementary Table 1). As recipient, we used DH10B cells with plasmid p*RFP* (Supplementary Table 1), which leads to red colored colonies on plate. If pF transfers to the recipient strain, transconjugants carry antibiotic resistance markers from both pF and pRFP (Figure 3A). By making use of the differences in antibiotic resistance (Figure 3B), we plated selectively for either recipient (Kanamycin) or transconjugant (Kanamycin and Tetracycline) colonies to assess the efficiency of conjugation (transconjugants per recipient). Colony counts for the transconjugants were on average three times lower than those of the recipients, yielding a conjugation efficiency of 0.35 transconjugants per recipient (T/R) (Figure 3C). Notably, we observed that the recipient plates (containing both the recipients and transconjugants) contained two classes of colonies: those that were either clearly red (as was observed earlier for DH10B with pRFP) and those that displayed a fainter red coloration. The transconjugant plates contained only faintly red colonies, suggesting that entry of pF into cells carrying pRFP represses mRFP protein production and/or maturation (Figure 3A). The ratio of faintly red (as opposed to clearly red) colonies on the recipient plates was 0.26 on average, quite close to the conjugation efficiency determined by selective plating (Figure 3C). In fact, we found that the fraction of faintly red colonies was more consistent between plates compared to the method of selective plating, likely because it does not require comparison of multiple plates. Through selective plating and colony PCR, we confirmed that the faintly red colonies were transconjugants (data not shown). It is concluded that DH10B is a suitable strain, both as a donor and as a recipient of the F plasmid and that pF inhibits red coloring of pRFP-carrying cells. The effect of pF on the color of colonies can therefore be used as an accurate and efficient method to measure the conjugation efficiency.

**Figure 3.**
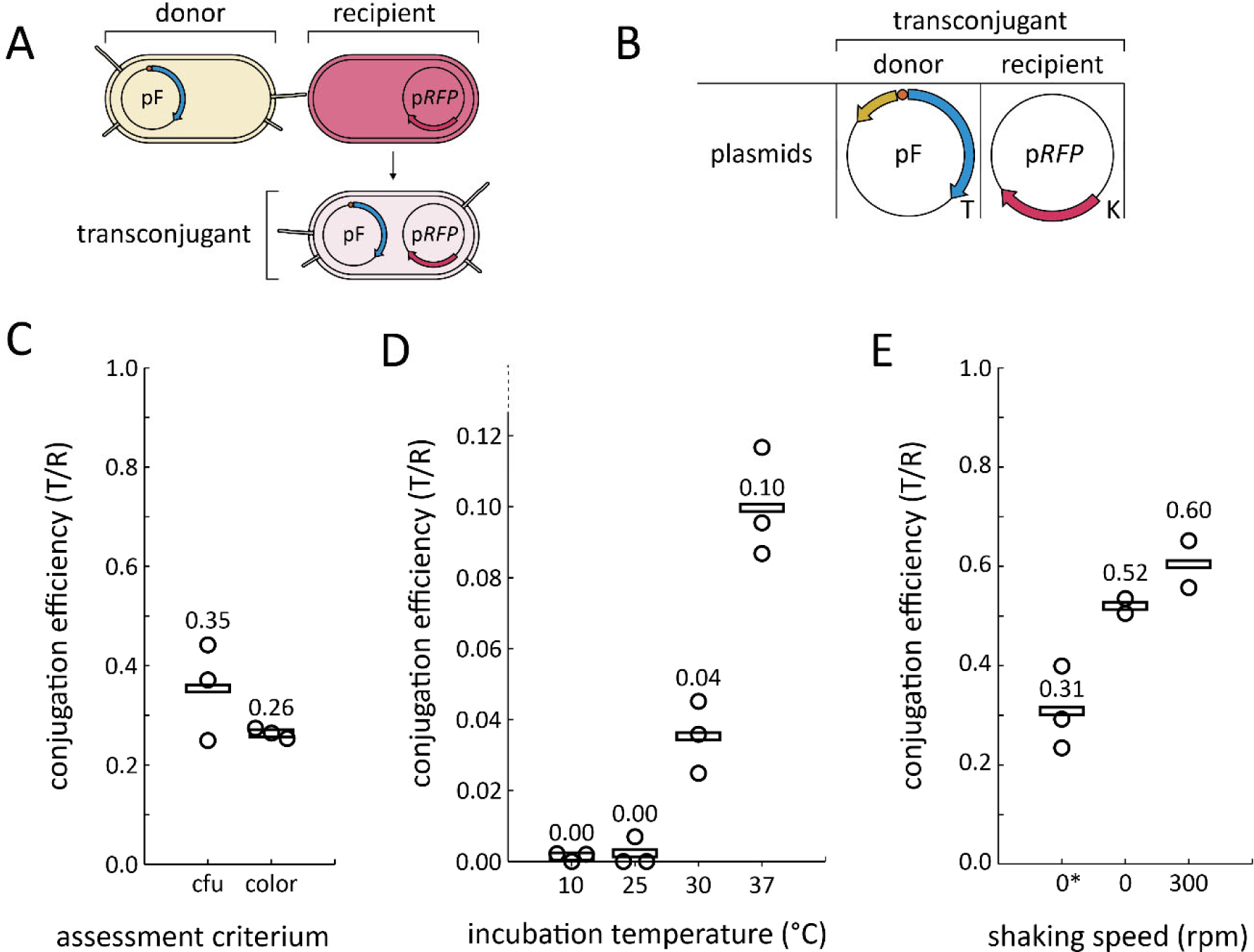
Conjugation by the F plasmid. For charts, individual data points are shown as small circles and the mean is indicated with a horizontal bar. **(A)** Schematic representation of the mating experiments for which data is shown in panels C, D, and E. Pili are shown to indicate that the donor and transconjugants would be able to initiate conjugation. The colors of the cells in the image are representative of the colors that were observed for colonies of these types of cells on plates. **(B)** The plasmids used for the mating experiments for which data is shown in panels C, D, and E. The names of the plasmids (Supplementary Table 1) are displayed in the center of each plasmid map, with the antibiotic resistance indicated on their bottom right. T: Tetracycline, K: Kanamycin. **(C)** Conjugation efficiency (transconjugants / recipients) as determined by two different methods: (i) comparing colony counts on plates with different antibiotics (cfu), and (ii) comparing the number of colonies on the recipient plates that were clearly colored red, or faintly red (color). Three biological replicates were used. **(D)** Conjugation efficiency at different temperatures. Three biological replicates were used. **(E)** Conjugation efficiency at different incubation conditions and shaking speeds. The asterisk indicates incubation in the incubator. The samples without an asterisk were incubated in the thermomixer. Three biological replicates were used for the ‘0*’ condition and two biological replicates were used for the other conditions.

We then investigated which conditions would be suitable for conjugation of the F plasmid in this setup. To that end, we conducted similar mating reactions at different temperatures and shaking conditions. We now exploited the pF-mediated inhibition of red coloring to measure the conjugation efficiencies in the different conditions. Of the tested temperatures, 37⁰C resulted in the most efficient conjugation (Figure 3D). Overall, conjugation efficiencies were much lower than those observed earlier (Figure 3C), likely because of differences in incubation, or starting concentrations of cells (further addressed in the Discussion section). In terms of shaking during the mating, we observed that 300 rpm shaking in a thermomixer can be used without negatively affecting conjugation efficiency (Figure 3E). It also seems that (regardless of shaking) samples in the thermomixer resulted in higher conjugation efficiencies, possibly because of more rapid heating of the tubes in the thermomixer.

### Plasmid spread through sequential conjugations

After conjugation, the resulting transconjugants should, in time, also be able to act as donors in secondary mating reactions. Sequential conjugation is a key requirement of the envisioned SCALE platform. We therefore tested sequential conjugations of the F plasmid and assessed whether these allow the F plasmid to spread throughout a bacterial population. For this, we used donor cells with both pF and pRFP and recipients with only pRFP (Figure 4AB). We used varying amounts of donor cells to inoculate cultures with 0%, 1%, 10%, or 100% of the starting cells being donors (carrying pF, Figure 4C). We then incubated these cultures in the thermomixer at 37⁰C with intermittent shaking, to allow mixing of the cells while not continuously disrupting pili and mating aggregates. We diluted the cultures daily in fresh medium and plated samples from each day to assess the presence of pF by the decreased red coloring of colonies. We did not differentiate between initial donors and cells that obtained pF through conjugation. As expected, the conditions that started with 0% or 100% donors were stable over time. In the intermediate conditions however, we observed that pF could spread throughout the populations (Figure 2D). In three days, pF had spread from 1% of cells to 100% of cells. This shows that the F plasmid can spread throughout a bacterial population through sequential conjugations in the tested conditions.

**Figure 4.**
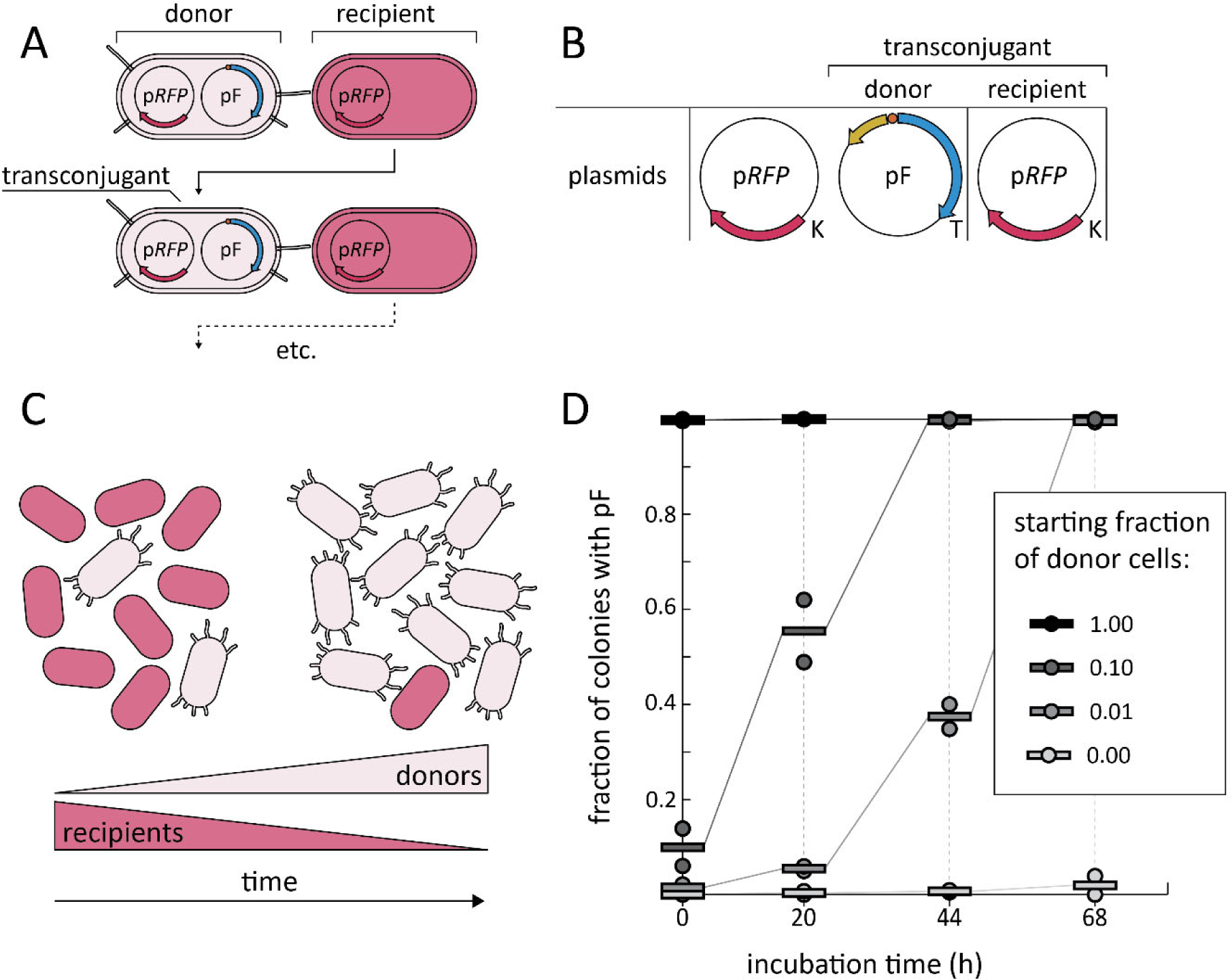
Plasmid spread through sequential conjugations. **(A)** Schematic representation of the mating experiments for which data is shown in panel D. Transconjugants are expected to become donors over time, allowing sequential conjugation. The colors of the cells in the image are representative of the colors that were observed for colonies of these types of cells on plates. **(B)** The plasmids used for the mating experiments for which data is shown in panel D. The names of the plasmids (Supplementary Table 1) are displayed in the center of each plasmid map, with the antibiotic resistance indicated on their bottom right. K: Kanamycin, T: Tetracycline. **(C)** Schematic representation of the expected shift in the numbers of donors and recipients over time. Donors: light pink with pili, recipients: bright red and smooth. **(D)** The fraction of colonies containing plasmid pF over time, when starting with different donor fractions (grayscales, see legend). Individual data points are shown as small circles and the mean is indicated with a horizontal bar. Two biological replicates were used.

### Transfer of a plasmid as facilitated by another

In the SCALE approach, mutations should not be accumulated in the transfer genes. Therefore, the GOI would ideally be expressed from a separate *oriT*-containing plasmid. Conjugation of the GOI-plasmid would then be facilitated by a different plasmid without *oriT*, expressing the transfer genes from the F plasmid. To confirm that conjugation of a secondary plasmid can be facilitated, we tested conjugation from donors carrying pF and p*RFP_oriT* (Supplementary Table 1) to recipients carrying p*GFP*, which carries a *GFP* gene and allows recipient-specific antibiotic selection. We obtained transconjugants and concluded that indeed the F plasmid can facilitate transfer of other plasmids carrying the corresponding *oriT* (data not shown). Notably, we again observed two types of transconjugants: those that were brightly red, and those that were faintly red (Supplementary Figure 1). This suggests that pF facilitates transfer of secondary plasmids (resulting in red colored transconjugant colonies), but also transfers itself to some of the transconjugants (faintly colored transconjugant colonies).

### Inhibition of conjugation by FinO

As discussed earlier, we found that selective plating can be used to assess conjugation, but that for exact quantification, other methods (such as loss of red color) might be more consistent across replicates. To allow for discrimination of small differences in conjugation efficiency, even more accurate and precise methods would be required. To this end, we investigated whether flow cytometry could be used to quantify conjugation efficiency.

In the envisioned SCALE platform (Figure 1), the FinO protein would be expressed from a separate plasmid. This *finO*-plasmid should, together with the FinP antisense RNA (expressed from pF), prevent conjugation until the *finO*-plasmid is cleaved and lost. In pF, the *finO* gene is naturally interrupted by insertion element IS3, rendering it unable to inhibit conjugation. To verify that FinO expressed from a separate plasmid could be used to prevent conjugative spread, we conducted conjugation experiments with donors containing pF and a reporter plasmid containing *oriT* and RFP, p*RFP_oriT* (Figure 5A, Supplementary Table 1). In two of the tested conditions, the donors were also supplied with a third plasmid: either p*finO* (which carries a reconstituted *finO* gene) or p*finO*_int (carrying the natural variant of the *finO* gene that is interrupted by IS3) (Figure 5A). The recipient cells all carried p*GFPuv* (Supplementary Table 1), which contains a *GFPuv* gene which can be used to visualize recipient cells (including transconjugants) during flow cytometry.

**Figure 5.**
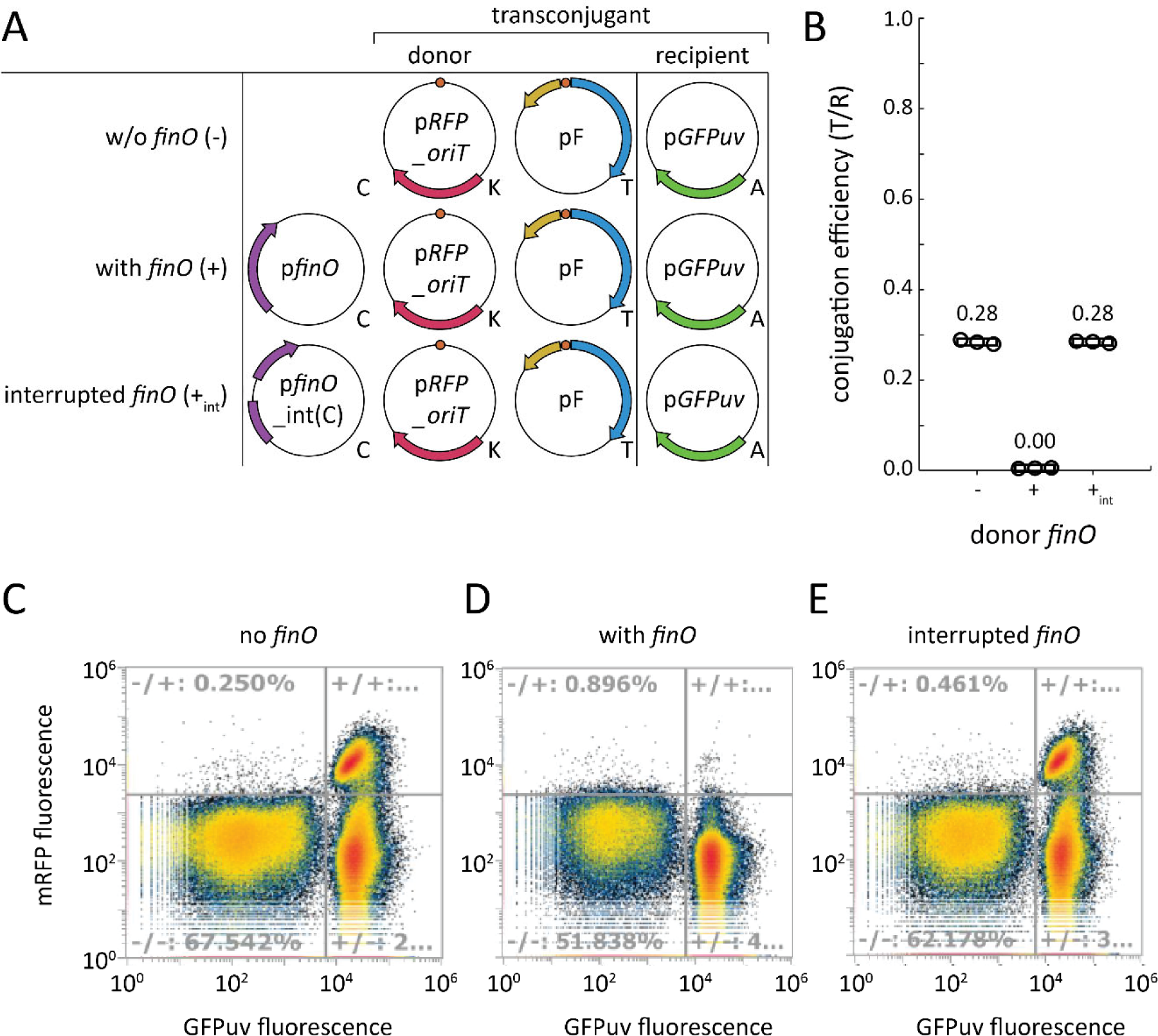
Prevention of conjugation by FinO. **(A)** The plasmids used for the mating experiments for which data is shown in panels B, C, D, and E. The names of the plasmids (Supplementary Table 1) are displayed in the center of each plasmid map, with the antibiotic resistance indicated on their bottom right. C: Chloramphenicol, K: Kanamycin, T: Tetracycline, A: Ampicillin. **(B)** Conjugation efficiency in the absence (-) or presence of the gene *finO*, either reconstituted (+) or interrupted by IS3 (+_int_). Conjugation efficiency was based on the data shown in panels C, D, and E. Conjugation efficiencies were inferred from the number of events in quadrants of mRFP and GFPuv fluorescence intensity. Three technical replicates were used. **(CDE)** Density maps of flow cytometry recorded events, with event density increasing from blue, to green, to yellow, to red. The logarithmic y-axes display the measured fluorescence intensity of RFP, while the x-axes show the same for GFPuv.

We found that the GFPuv signal could be used to visualize the recipient cells (including transconjugants), and RFP signal could be used to visualize the cells with p*RFP_oriT* (donors and transconjugants) (Figure 5A) with minimal interference from each other. These signals were used to group cells into donors, recipients, and transconjugants (Figure 5C-E). In the bottom left quadrants (low GFPuv, low RFP) we assume to observe donor cells. The RFP signal is likely low for the donor cells because of the presence of pF, as discussed earlier. In the bottom right quadrant (high GFPuv, low RFP), we see recipient cells that did not receive the p*GFPuv* plasmid. On the top right (high GFPuv, high RFP), transconjugants are seen. The number of gated events for each group was used to infer conjugation efficiencies (transconjugants per recipient) for each donor type. We observed that donors without *finO*, or with an interrupted *finO* gene (p*finO_*int(C)) showed efficient conjugation, whereas conjugation efficiency in the presence of active *finO* (p*finO*) was successfully diminished (Figure 5B). Across technical replicates, we observed little variation. This indicates that flow cytometry is a precise method for assessing conjugation efficiency, likely because of the high throughput of this approach.

### Streamlining expression of genes required for conjugation

In SCALE, instead of having the entire F plasmid present, it would probably be better in terms of efficiency and stability, to integrate the genes from the F plasmid that are required for conjugation on the bacterial chromosome. To identify which genes are required, we assembled a 53kb streamlined version of the F plasmid (∼100kb), pStream (Figure 1B), which contains the tra-region, the leader, *oriT*, and the elements needed for replication and stable maintenance (Supplementary Table 1). We tested conjugation of donors with pStream to recipients with p*RFP* and conducted similar experiments in parallel with donors containing pF as a positive control (Figure 6A). Selective plating of transconjugants showed that pStream had a conjugation efficiency similar to pF (Figure 6B). This shows that the tra-region, the leader, and *ori*T are sufficient for efficient conjugation by the F plasmid. Strikingly, transconjugants with pStream did not display the reduction in red color which had been observed previously for pF.

**Figure 6.**
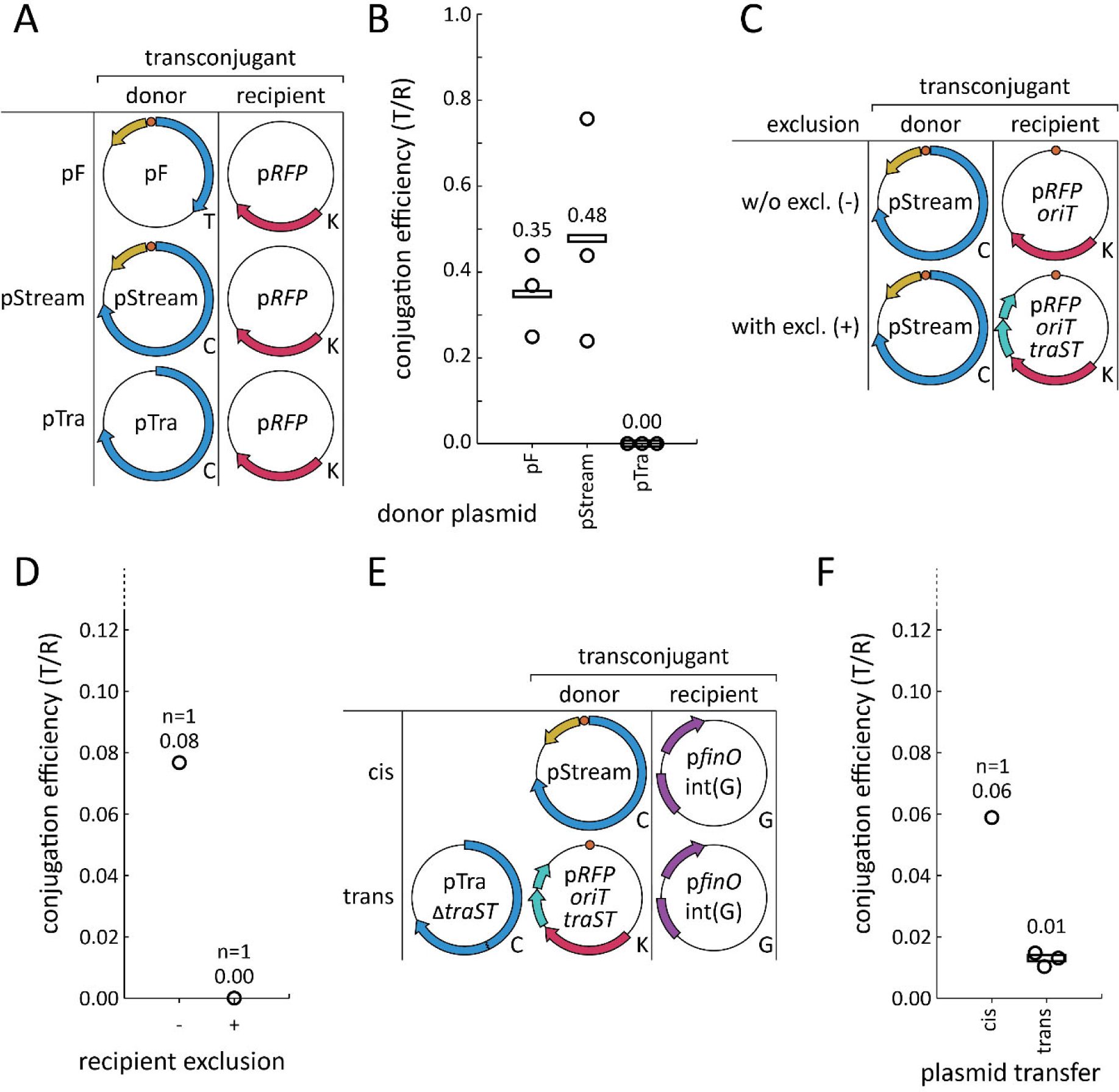
Conjugation as facilitated by the tra-region. For charts, individual data points are shown as small circles and the mean is indicated with a horizontal bar. **(A)** The plasmids used for the mating experiments for which data is shown in panels B. The names of the plasmids (Supplementary Table 1) are displayed in the center of each plasmid map, with the antibiotic resistance indicated on their bottom right. **(B)** Conjugation efficiency (transconjugants / recipients) for donors with either pF, pStream, or pTra. Three biological replicates were used. **(C)** The plasmids used for the mating experiments for which data is shown in panels D. **(D)** Conjugation efficiency (transconjugants / recipients) to recipients without (-) or with (+) *traS* and *traT* genes. **(E)** The plasmids used for the mating experiments for which data is shown in panels F. **(F)** Conjugation efficiency (transconjugants / recipients) for donors with either pStream (cis) or both pTra_Δ*traST* and p*RFP_oriT_traST* (trans). T: Tetracycline, K: Kanamycin, C: Chloramphenicol, G: Gentamycin.

After establishing that the remaining F plasmid elements on pStream are sufficient for efficient conjugation, we next assembled plasmid pTra which contains the tra-region, but neither the *oriT* nor the leader sequence from the F plasmid (Figure 1C, Supplementary Table 1). pTra was intended to facilitate conjugation of a secondary plasmid without transfer of pTra itself. We indeed found, through conjugation (Figure 6A) and selective plating, that the absence of *oriT* and the leader region prevented conjugation of the pTra plasmid (Figure 6B).

In the envisioned SCALE platform, the plasmid with the GOI is transferred to cells containing the genes needed for conjugation. However, these cells also need to be able to act as recipients to obtain the GOI- containing plasmid in the first place. If pTra would be used, the *traS* and *traT* genes would prevent such a mating through the mechanisms of surface- and entry-exclusion. To address this limitation of pTra, we constructed pTra_Δ*traST* (Figure 1D, Supplementary Table 1) in which *traS* and *traT* are knocked out.

The *traS* and *traT* genes of the F plasmid prevent redundant mating reactions. Complete absence of *traS* and *traT* might therefore increase the burden on the bacterial population due to uncontrolled conjugations. To prevent this, we constructed a new version of p*RFP_oriT*, p*RFP_oriT_traST* (Supplementary Table 1) that contains the *traS* and *traT* genes. Because p*RFP_oriT_traST* is the plasmid that is transferred, *traS and traT* on this plasmid should prevent superfluous conjugation attempts in transconjugants. When either p*RFP_oriT* or p*RFP_oriT_traST* was used as a recipient with donors containing pStream (Figure 6C), we saw that indeed only conjugation to p*RFP_oriT* was successful (Figure 6D), showing that *traS* and *traT* from p*RFP_oriT_traST* prevent conjugation. We then tested whether pTra_Δ*traST* could be used to transfer p*RFP_oriT_traST* in trans (Figure 6E), and observed transconjugants, but at very low conjugation efficiency compared to self-transfer of pStream (Figure 6F).

To improve the inefficient conjugation with pTra_Δ*traST*, we constructed p*RFP_oriT_traST*_*slt* (Supplementary Table 1), which is similar to p*RFP_oriT_traST*, but includes the *slt* (soluble lytic transglycosylase) gene from the F plasmid leader, which is involved in local peptidoglycan degradation (69, 70) and had been reported to increase conjugation efficiency (71). When we quantitively assessed conjugation efficiencies (Figure 7A), we found that the *slt* gene in p*RFP_oriT_traST*_*slt* indeed improved conjugation efficiency substantially compared to p*RFP_oriT_traST* (Figure 7B). This confirmed that the *slt* gene is an important contributor to F plasmid-mediated conjugation.

**Figure 7.**
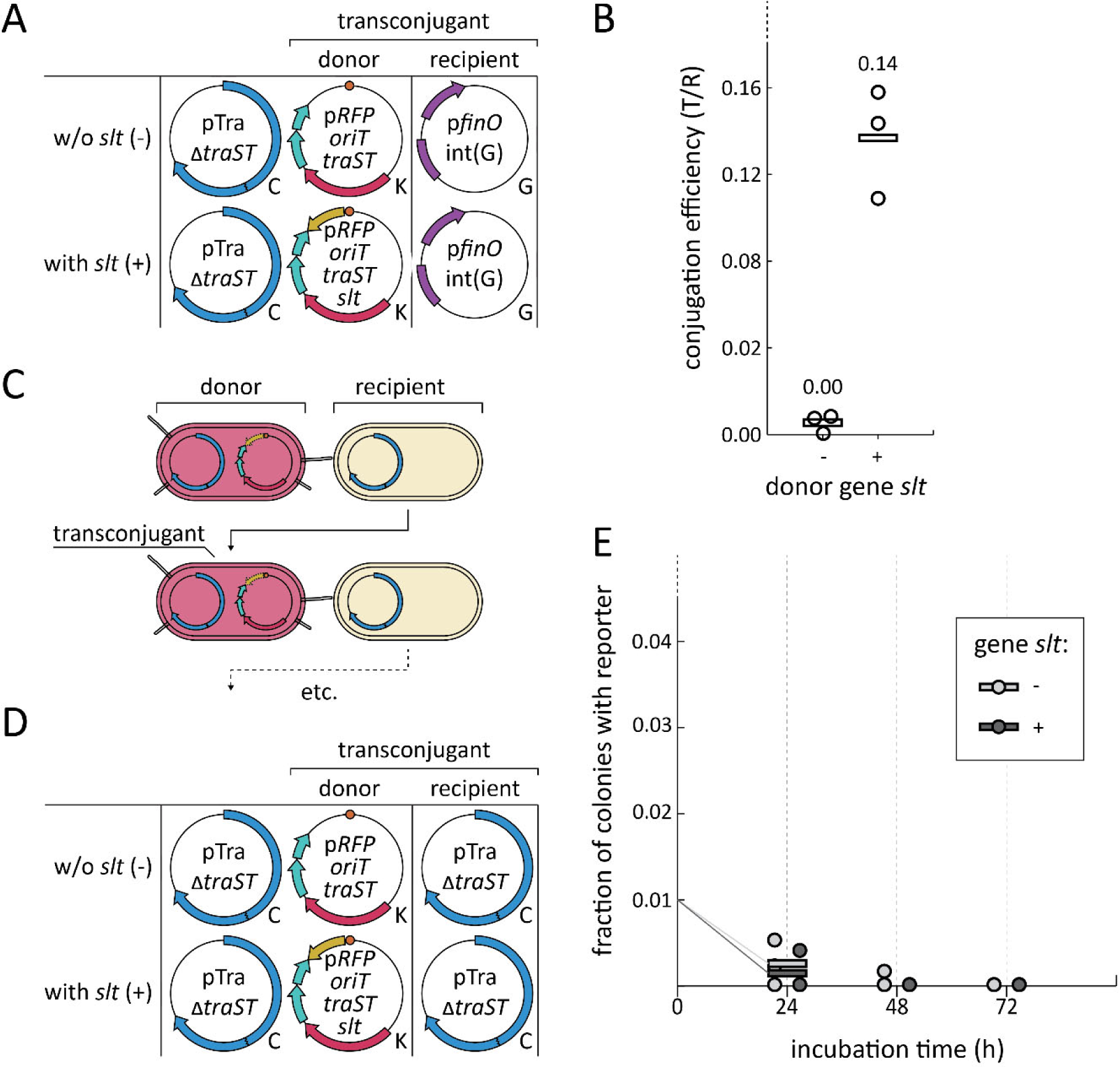
The effect of gene *slt* on conjugation and conjugative plasmid spreading. For charts, individual data points are shown as small circles and the mean is indicated with a horizontal bar. **(A)** The plasmids used for the mating experiments for which data is shown in panels B. The names of the plasmids (Supplementary Table 1) are displayed in the center of each plasmid map, with the antibiotic resistance indicated on their bottom right. **(B)** Conjugation efficiency (transconjugants / recipients) for donors either without (-) or with (+) the *slt* gene. Three biological replicates were used. **(C)** Schematic representation of the mating experiments for which data is shown in panel E. Transconjugants are expected to become donors over time, allowing sequential conjugation. The colors of the cells in the image are representative of the colors that were observed for colonies of these types of cells on plates. **(D)** The plasmids used for the mating experiments for which data is shown in panels E. **(E)** The fraction of colonies containing plasmid p*RFP_oriT_traST* or p*RFP_oriT_traST_slt* (donors, see legend) over time. Three biological replicates were used. C: Chloramphenicol, K: Kanamycin, G: Gentamycin.

We then tested if, given the now improved conjugation efficiency, pTra_Δ*traST* could facilitate sequential conjugations of p*RFP_oriT_traST*_*slt* to spread it through a bacterial population (Figure 7C). To this end, we used mating reactions where all cells contained pTra_Δ*traST* and 1% of cells (the donors) also contained p*RFP_oriT_traST* or p*RFP_oriT_traST*_*slt* (Figure 7D). We found that neither p*RFP_oriT_traST* nor p*RFP_oriT_traST*_*slt* could spread through the population within three days (Figure 7E).

### Chromosomal integration of the transfer region

We hypothesized that spreading was unsuccessful because the transferred plasmids lack elements from F that are needed for sequential conjugations. Compared to pF and pStream (for which we have observed efficient spreading, data not shown for pStream), plasmid p*RFP_oriT_traST* and p*RFP_oriT_traST_slt* lack the replication and maintenance factors of the F plasmid, as well as the leader region. Expression of the transfer region from the chromosome would allow us to use the F plasmid backbone (which can accommodate large pieces of DNA like the leader region) for the transferring reporter plasmid. To explore this option, we used the F plasmid backbone to construct p*lacZ_traST_slt_oriT*, which carries the *lacZ*, *traS, traT, slt,* and *oriT*. In addition, we constructed p*lacZ_traST_slt_oriT*, which is similar but lacks the *oriT*. We then attempted to integrate the tra-region from pTra_Δ*traST* into the *E. coli* chromosome (Figure 1D, E). To integrate such a large (32kb) DNA sequence using homologous recombination, we planned to first integrate a ‘landing site’ on the chromosome containing a sequence homologous to the tra-region of the F plasmid. In a second integration step, the tra-region would then be inserted into the landing site by homologous recombination.

We first attempted to integrate the landing site at the *attB* locus, the integration site for phage lambda using homologous recombination and CRISPR-Cas counterselection (72), but we were not able to identify any successful integration at that site. We then attempted to integrate the landing site into three other genomic regions (73, 74). In all three cases we did obtain successful integration of the landing site. We then tried to integrate the tra-region from pTra_Δ*traST* into these three different landing sites. For one location, we succeeded as verified by sequencing (data not shown). Unfortunately, when we tested conjugation of p*RFP_oriT_traST_slt* and p*lacZ_traST_slt_oriT* from the resulting strain, we did not observe any transconjugants, as assessed by spot plating (Figure 8). This shows that either the integrated genes or not expressed properly, or that these genes are not sufficient to facilitate conjugation in trans.

**Figure 8.**
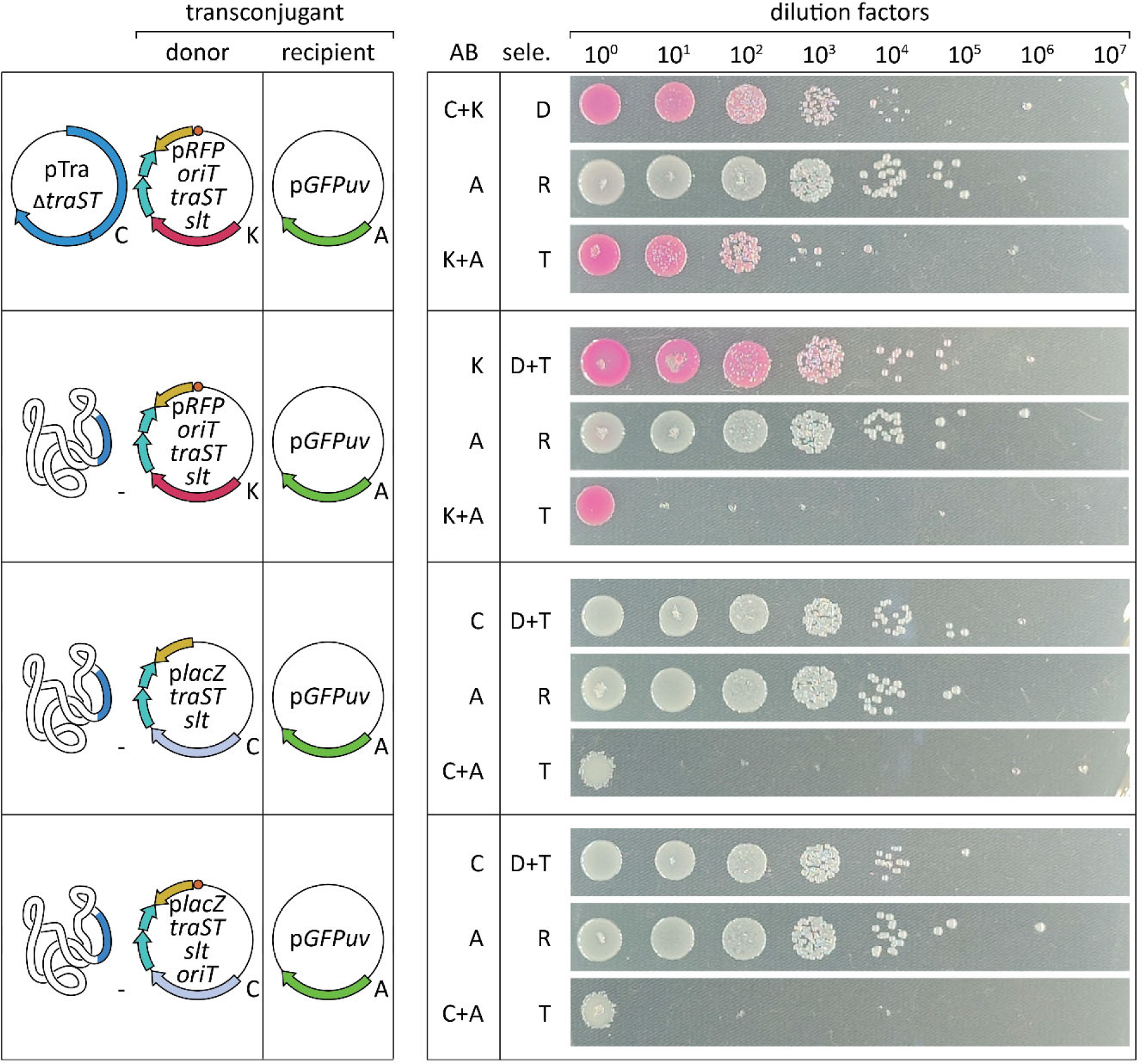
Spot plating assay of conjugation facilitated by chromosome-integrated tra-region. From left to right, the figure shows (i) The plasmids used for the mating experiments. The names of the plasmids (Supplementary Table 1) are displayed in the center of each plasmid map, with the antibiotic resistance indicated on their bottom right. (ii) The antibiotics of the plates (AB). (iii) Which types of cells (donors, recipients, and/or transconjugants) should growth on the plates (sele.). (iiii) Photos of the spot plating plates. Spots were pipetted from serial dilutions (dilution factors indicated on top) after the mating reactions. Three biological replicates were used, and pictures are representative photos from the triplicates.

## Discussion

We have investigated whether F plasmid conjugation could be used for a laboratory evolution platform for DNA endonucleases and other DNA-affecting proteins. Initially, we tested the effect of different temperatures and shaking conditions on the efficiency of conjugation. Out of the tested conditions, 37°C was optimal (Figure 3D), mirroring the ideal growth temperature of *E. coli*. This is in line with previous reports that conjugation is tightly connected to bacterial growth (51). Slow shaking in a thermomixer was well-tolerated (Figure 3E), which might be advantageous to guarantee mixing of donor and recipient cells over longer incubation times. The F plasmid can transfer in liquid mating reactions due to mating pair stabilization (53). This is expected to be advantageous for laboratory evolution, as cells can be mixed to allow efficient conjugation over longer times. Indeed, we have shown that in laboratory conditions, we could spread the F plasmid through a bacterial culture (Figure 4D). In addition, we found that conjugation and spreading could be prevented by introduction of the reconstituted *finO* gene (Figure 5B, D). To reconstitute the *finO* gene, we constructed a version lacking the IS3 element, restoring the reading frame. As expected, the interrupted version of *finO* did inhibit conjugation (Figure 5B, E).

We noted that conjugation efficiencies can differ substantially, even between similar experiments (Figure 2B, 2C). This is likely due to small differences in the execution of the experiments such as differences in starting OD_600_ or the exact duration and temperature of incubation (75). This highlights the importance of only comparing conditions that were included in the same experiment, to minimize these differences. Because of this, we have refrained from quantitatively comparing results between different experiments.

Initially, the *slt* gene (also called *ygfA*, gene *19*, gene *X*, ORF169) from the F plasmid had not been included in conjugations with pTra and pTra_Δ*traST* because all elements needed for conjugation are supposedly located in the tra-region (47), which does not include the *slt* gene (Figure 1A,B). However, previous work by others has indicated that an orthologous *slt* from F-like plasmid R1 does contribute to increased conjugation efficiency (71). This is likely due to local degradation of the peptidoglycan layer, allowing construction of pili or mating pores on the cell surface (69, 70). Indeed, inclusion of *slt* in our experiments improved conjugation efficiency by one order of magnitude (Figure 3D), consistent with what had been observed by others.

Although we made progress in evaluating the potential of F plasmid conjugation for laboratory evolution, we have not been able to fully establish the envisioned platform. The main bottleneck in our studies was that we were not able to use transfer genes to facilitate conjugative plasmid spread of a separate plasmid (Figure 7E). The most probable explanation for this is that the used reporter plasmids do not carry the leader region from the F plasmid, which is likely important for sequential conjugations (55). The leader region is quite large (around 13kb), making it difficult to incorporate it into p*RFP_oriT_slt*. The F plasmid backbone, however, would be able to accommodate the leader region. Therefore, if we manage to functionally integrate the tra-region into the bacterial chromosome, we could use a reporter plasmid with *oriT*, the F plasmid backbone, and the full leader region (as well as *traS* and *traT*).

Alternatively, *traS* and *traT* might explain the failed sequential conjugations with the pTra_Δ*traST* construct. Both *traS* and *traT* have been knocked out to construct pTra_Δ*traST*, but not completely (see materials and methods). However, it seems highly unlikely that any activity remains because there is only 91bp and 70bp left of the coding sequence of *traS* and *traT* respectively (originally, *traS* and *traT* coding sequences are 522 and 735bp long respectively).

Another advantage of using the F plasmid backbone for the GOI-plasmid is the backbone’s low copy number (2 copies per chromosome (76)). This would prevent confounding effects of having many different GOI variants within a single cell (where variant A might pass the selection criteria, causing variant B to be transferred to the next evolutionary cycle). So far, we have indeed attempted to integrate the tra-regions from pTra_Δ*traST* into the chromosome of DH10B, but the resulting strain could not facilitate conjugation of secondary plasmids (Figure 4). We have sequence-verified correct integration through whole genome sequencing.

In addition, we wonder whether the presence of the conjugative genes in the chromosome could be toxic or pose a large burden to the cells. This would be somewhat surprising because we know that the same genes can be tolerated when expressed from a plasmid (pTra_Δ*traST*). In addition, high-frequency recombination (Hfr) strains resulting from integration of the F plasmid in the bacterial chromosome are both viable and show active conjugation. In our attempts, *oriT* is purposefully not integrated on the chromosome to prevent nicking and attempts of transferring the chromosome itself, as happens for Hfr strains. Still, we would like to exclude the possibility of toxicity interfering with the chromosomal integration, by first introducing a plasmid carrying *finO*, which inhibits conjugation. In this strain we would then again attempt integration of the tra-region. It would be best if expression of *finO* can be inhibited, or if the plasmid carrying *finO* is easy to cure (through targeting with Cas9 for instance). If the integration of the tra-region is successful, but *finO* repression or curing is lethal, we would conclude that the integrated tra-region is indeed toxic or poses a large cellular burden.

If functional integration of the tra-region is achieved, the next step is testing whether the resulting strain allows spreading of an *oriT*-containing plasmid (p*lacZ_traST_slt_oriT*) through sequential conjugations. If conjugative spread is observed, the next step is to test whether *finO* cleavage can be used to selectively spread plasmids with functional nucleases. To that end, two adaptations of p*lacZ_traST_slt_oriT* should be tested: one expressing a functional Cas9, and another with a catalytically inactive version of Cas9. These plasmids should then be challenged to spread through a population of cells that contain both the chromosomal integration of the tra-region, and a *finO* plasmid targeted by Cas9. The plasmids encoding active Cas9, should lead to cleavage and subsequent degradation of the *finO* plasmid. This in turn would relieve the repression of conjugation, allowing the Cas9-carrying plasmid to transfer to other cells.

Finally, to test enrichment of active variants, donor cells containing either of the two Cas9 plasmids (active or inactive) should be mixed in different ratios. Starting from a ratio of 1:100 active versus inactive Cas9, selective conjugative spread of plasmids with the active variant should enrich for these plasmids. After three days, the resulting culture should be plated, and sequencing of individual colonies should show whether the plasmid encoding active Cas9 is indeed enriched. If this point is reached, we would conclude that F plasmid conjugation is useful as a method to select DNA affecting proteins.

If chromosomal integration of the tra-region fails, SCALE could theoretically be realized with just plasmids. The first plasmid-only alternative (alternative 1) would be to express the GOI from pStream (Figure 9A, B). The major limitation of this approach is that the mutations on pStream would propagate to recipient cells. In the second alternative (alternative 2), the GOI would be expressed from a separate plasmid that also contains *oriT* (Figure 9A). Conjugative transfer of this ‘p*GOI_oriT*’ plasmid would be facilitated by pStream, which also transfers itself. The TraS and TraT proteins encoded on pStream would prevent conjugation only once their concentrations are high enough, potentially allowing p*GOI_oriT* to be transferred as well (Figure 9C). We have observed p*RFP_oriT* being transferred along with pF (Figure 5C), showing that conjugative transfer of p*GOI_oriT* would be possible in this setup. However, it remains to be seen whether this approach is suitable for sequential conjugations too. For both plasmid-only alternatives, new plasmid would have to be constructed with a unique antibiotic resistance on each.

**Figure 9.**
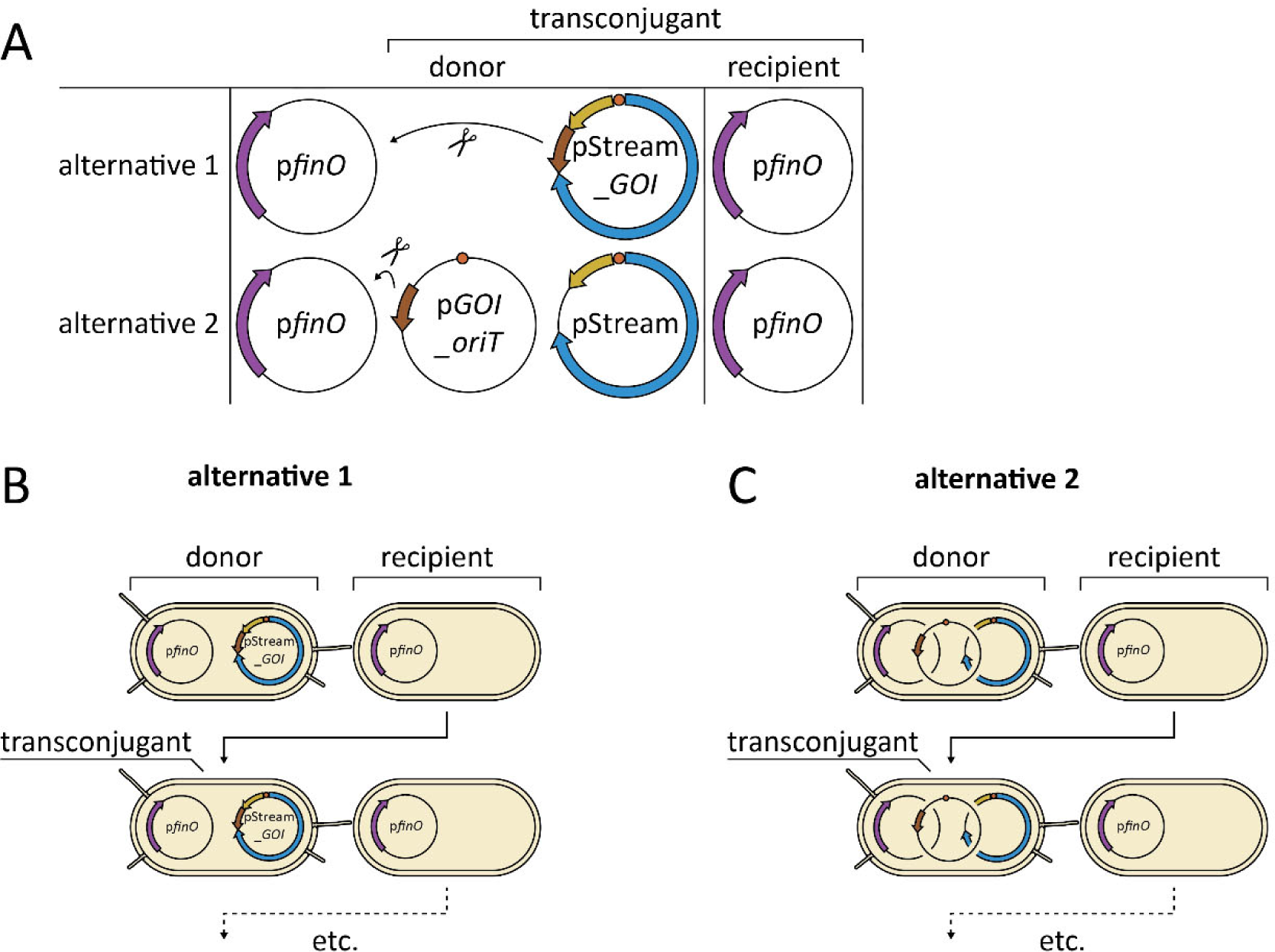
Plasmid-only alternatives to chromosomal integration of tra-region. **(A)** The plasmids that would be used in the proposed plasmid-only alternatives. Arrows with a pair of scissors indicate that the GOI would result in cleavage of the p*finO* plasmid in order to allow conjugative transfer. **(BC)** Schematic representation of the mating reactions that would take place for the proposed plasmid-only alternatives. Transconjugants are expected to become donors over time, allowing sequential conjugation.

A useful laboratory evolution platform requires not just a selection method, but mutagenesis of the GOI as well. Although spontaneous mutations would occur within the GOI, the rate is likely too low to allow meaningful laboratory evolution on a short timescale. Therefore, it would be desirable to combine the conjugative selection system with a fully *in vivo* mutagenesis approach. The most straightforward solution would be to use the mutagenesis plasmid MP6 (39) which increases the overall rate with which mutations arise within bacterial cells. Because only the GOI-plasmid is transferred, and because new recipients can be supplied in each passage, mutations occurring outside the GOI-plasmid should not interfere with the selection approach.

Another mutagenesis option we have considered is to use an error prone DNA polymerase I (77, 78) in conjunction with a GOI-plasmid with a ColE1 origin of replication. DNA polymerase I starts replication ColE1 plasmids (79, 80), but is replaced by polymerase III after approximately 170-250 nucleotides (80). Mutation frequencies from error prone polymerase I were indeed highest close to the origin of replication but remained relatively high for at least 3kb downstream from there (77), compared to the wild-type polymerase. Because Cas nucleases like Cas9 and Cas12a are quite large proteins, mutagenesis of long DNA sequences is important.

Yet another method for mutagenesis is the use of a T7 RNA polymerase fused to a cytidine deaminase (81, 82). The GOI would then be expressed under a T7 promoter, causing transcription by the T7 RNA polymerase. The T7 polymerase thereby recruits the cytidine deaminase to the GOI, leading C-G base pairs to be converted to T-A base pairs. If the promoter is also included downstream from the GOI as reverse complement, G-C base pairs can also effectively be converted to A-T pairs. This method should have little effect on most DNA in the cell (as mutagenesis is restricted to T7 promoter-flanked sequences), specifically mutating the GOI, making it a suitable mutagenesis strategy in combination with the above described ‘alternative 1’ selection approach (Figure 9A). On the other hand, C to T (and G to A) conversions constitute a limited mutagenesis spectrum, excluding the other conversions, insertions, and deletions. Perhaps other conversions could be made possible by constructing an adenine deaminase fusion to T7 RNA polymerase as well.

Overall, we have proposed SCALE, a laboratory evolution platform for DNA cleaving proteins, based on plasmid conjugation. We have established sequential conjugation, a streamlined conjugative plasmid, as well as a system where conjugation of a plasmid is facilitated by conjugative plasmids expressed from another plasmid. Unfortunately, we were not able to conclusively test whether SCALE would work to enrich DNA targeting proteins like Cas9. A functional integration of the F plasmid transfer region into the bacterial chromosome would likely be an important step for future studies. Alternatively, plasmid-only approaches might also allow DNA-cleavage dependent transfer of the GOI between cells. If the envisioned laboratory evolution platform can be realized, it might aid applications of Cas nucleases by allowing easy fine-tuning toward their intended use. For instance, Cas nucleases could be evolved toward different PAMs, increased specificity, or reduced size. In addition, the approach described would – if functional – also be suitable for laboratory evolution of other proteins that change the DNA sequence, such as base editors and prime editors.

## Methods

### Bacterial strains and growth conditions

The F plasmid was obtained from *Escherichia coli* strain ER2738, provided by New England Biolabs (NEB). For all conjugation experiments, the *E. coli* strain DH10B was used both as donor and recipient. Cultures were grown in LB (10g/L NaCl, 10g/L Peptone, 5g/L Yeast extract) or on plates of LB agar (LB with 15g/L agarose), both containing the appropriate antibiotics. For antibiotics we have used Ampicillin (working concentration 100 µg/mL), Chloramphenicol (25 µg/mL), Spectinomycin (100 µg/mL), Kanamycin (50 µg/mL), Gentamicin (50 µg/mL), and Tetracycline (5 µg/mL). Typically, we grew DH10B overnight in 5mL LB in 50mL conical tubes at 37°C.

### Plasmid cloning

Plasmids below 10kb in size were constructed through PCRs and subsequent Hifi assembly (NEB E2621S), according to manufacturer’s protocol. Larger plasmids, specifically those derived from the F plasmid, were produced through a combination of restriction digestion and Hifi assembly with PCR amplified DNA. DNA of large, low copy plasmids was purified mostly using the Macherey Nagel NucleoBond Xtra BAC kit (product number 740436). pStream, pTra, and pTra_Δ*traST* were constructed using pBeloBAC11 (NEB) as plasmid backbone. To construct pTra_Δ*traST*, we used five BsaI restriction sites that are naturally present within the tra-region of the F plasmid, close to the *traS* and *traT* genes. We used Hifi assembly to reconstitute the genes surrounding *traS* and *traT* (*traG* and *traD*), using a second fragment created through overlap-extension PCR on these genes. The resulting pTra_Δ*traST* construct contains the first 91bp of *traS* and the last 70bp of *traT*, minimizing the chance of disrupting expression of other genes in the tra-region.

### Conjugative mating reactions

For conjugations, we followed a slightly adapted version of a referenced (75) protocol for liquid matings. We started 5mL overnight pre-cultures with appropriate antibiotics. After overnight incubation, these cultures were washed twice by centrifugation at 1200 rcf for 10min, then discarding the supernatant, and finally resuspending the bacterial pellet in 1mL fresh LB without antibiotics. We used 20µL of the resuspended cells to inoculate a culture of 5mL LB without antibiotics in a 50mL conical tube, which was then incubated at 37°C for 2h with vigorous shaking. After incubation, OD_600_ values of the cultures were measured and normalized through dilution with LB without antibiotics. We then mixed 200µL of donor cells with 200µL of recipient cells in a 2mL microcentrifuge tube and incubated at 37°C for 1h without shaking or shaking speeds up to 40rpm. To stop the conjugation, samples were vortexed for 1min or incubated at room temperature at 1000 rpm shaking for 10min.

### Conjugative spreading

To assess whether plasmid could spread throughout a population by sequential conjugations, we conducted spreading experiments. These experiments were prepared in the same way as regular matings, through overnight cultures, which were washed, and grown in LB without antibiotics. OD_600_ values were used to normalize the concentrations of cells in each culture and donor and recipients were mixed. In contrast to regular matings, donors and recipients were mixed in different ratios and diluted 100x before incubation (OD_600_ 0.02-0.04). We used 2mL microcentrifuge tubes with 200µL or 300µL total volume of cells and medium. These co-cultures were then incubated for a longer duration, up to multiple days in 2mL microcentrifuge tubes in the thermomixer (Eppendorf Thermomixer C) at 37⁰C with intermittent shaking at 1000 rpm for 15min, followed by 1h without shaking. The periods without shaking are included because vigorous shaking is known to disrupt the F pili and prevent conjugation. To stop the conjugation, samples were incubated at 37°C at 1000 rpm shaking for 15min in the thermomixer.

### Selective plating for quantifying conjugation

To assess conjugation efficiencies, we plated the samples on LB agar with selective antibiotics for either donors, recipients, or transconjugants. To this end, we used OD_600_ values of conjugation samples to estimate the number of colony forming units (cfu) per mL and used LB to dilute the samples to an estimated cfu/mL of 1000-3000. We then plated 75-100µL of the resulting dilution on LB agar with selective antibiotics and incubated at 37°C overnight. In general, the ratio between donors, recipients, and transconjugants was determined by counting colonies on plates with the corresponding antibiotics. In some cases, where this is mentioned in the text, the fractions fluorescent or colored colonies were used to infer the fractions of donors, recipients, or transconjugants. Here it is important to realize that recipients consist out of two types of cells: those recipients that obtained the donor plasmid (i.e., transconjugants), and those that did not. Transconjugant are thus always also recipients, but not the other way around.

### Spot plating for quantifying conjugation

Serial spot plating is an alternative way of selective plating, also using selective antibiotics to assess donors, recipients, and transconjugants. Instead of plating a single dilution on an LB agar plate, in spot plating we pipetted 3µL droplets of different concentrations on single plate. This was used to cover a wide range of cfu concentrations efficiently.

### Flow cytometry for quantifying conjugation

To assess conjugation efficiency with flow cytometry, mating reactions were performed as described above. After 1h of mating, the reactions were incubated at 4°C for 3h, intended to mature RFP and GFPuv for fluorescence, without allowing growth or additional conjugation. Flow cytometry was performed on the Attune NxT (software version 3.1.1). Side scatter was adjusted to exclude non-cellular events. Fluorescence was measured with a VL2 and YL2 filter, for GFPuv and RFP fluorescence respectively. The number of donors, recipients, or transconjugants was inferred from the fraction of cellular event within the corresponding quadrant of GFPuv and RFP intensity. The used voltage for forward scatter, side scatter, GFPuv fluorescence and RFP fluorescence were 350, 350, 300, 400V respectively. For each analysis, we aimed to obtain 10.000-20.000 events per second, using a flow of 25µL/min, by appropriately diluting the samples. Gating was set using cells with p*RFP* and cells with p*GFPuv* as standards.

### Chromosomal integration of tra-region

We tried to integrate the tra-region into the chromosome of DH10B in two steps, first creating a landing-site which could be used in the second step to integrate the tra-region itself. For these chromosomal integrations we used the referenced (72) method which uses a combination of lambda red homologous recombination and counterselection by Cas9. Originally, this system used two plasmids: pCas (which carries *cas9* and lambda red) and pTarget. We first used pTarget plasmids with guides (which target the chromosomal sequence where the integration would take place) and repair templates that contain regions homologous to the outer regions of the tra-region, flanked by homology arms to match the chromosomal region around the cleavage site of Cas9. The regions where we integrated the landing site were the “*ypjC_ileY* intergenic region” (73), “integration site 7” (74), and “safe site 9” (74). These sites were selected because they have shown decent levels of expression of integrated genes, they should not disrupt other *E. coli* genes, and because they have a suitable Cas9 protospacer. After integration of the landing site, we cured the first pTarget plasmids and introduced new pTarget plasmids without a repair template, only carrying guides that target the landing site DNA (but not the tra-region). In this last step, the homologous template is supplied from pTra_Δ*traST*. We used 500bp homology arms for integration of both the landing site and the tra-region.

## Acknowledgment

We thank Fernando de la Cruz, Nico Claassens, and Sjoerd Creutzburg for advice in the early stages of this project. Bram Sturkenboom was involved in the first attempts to integrate the transfer region of the F plasmid in the *E. coli* chromosome.

## Supplementary information

**Supplementary Figure 1.**
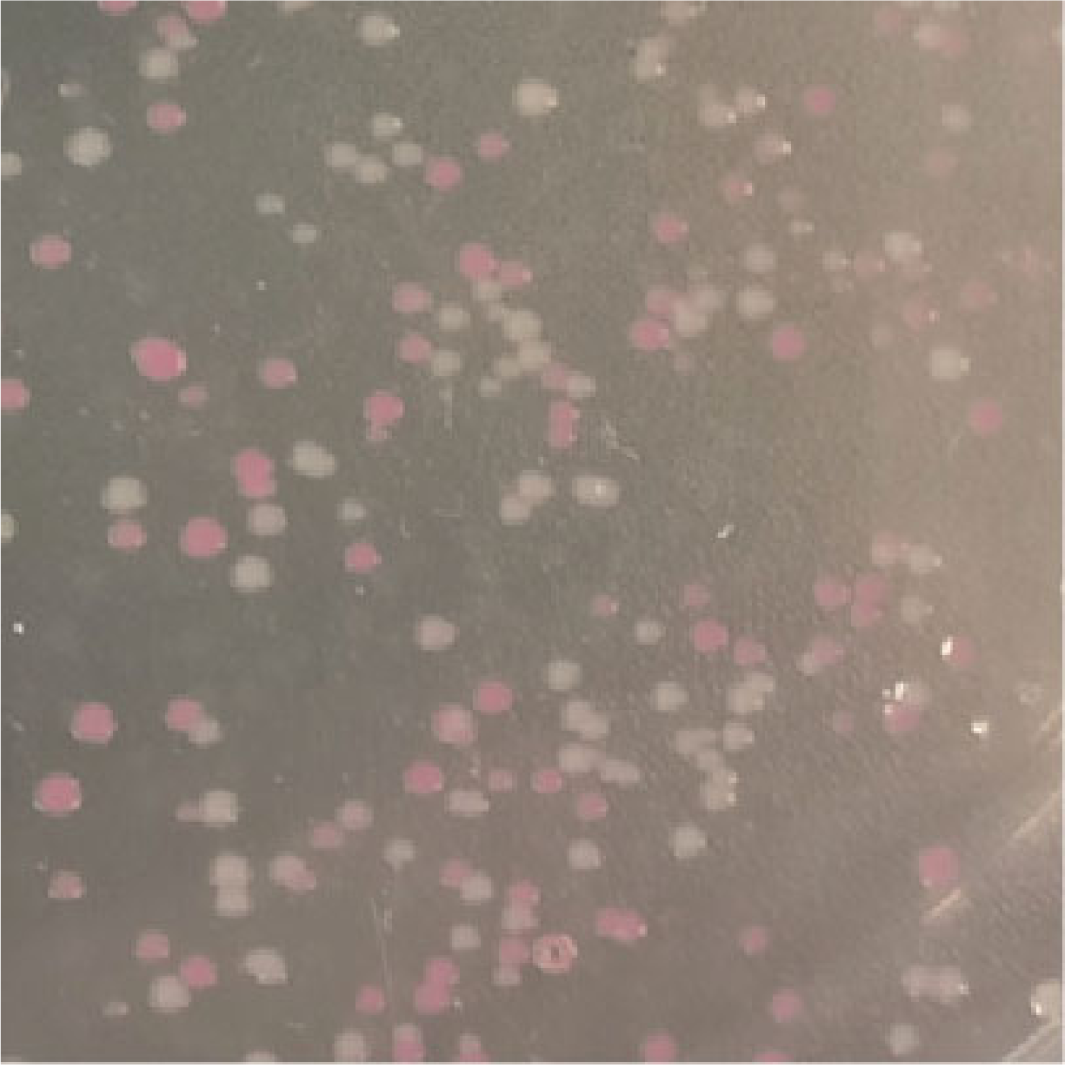
Different intensities of RFP depending on the presence of pF. Photo of a selective (Kanamycin and Ampicillin) LB agar plate for transconjugants of a mating of DH10B donors with pF and p*RFP_oriT*, with recipients carrying p*GFP*. The plasmids pF, p*RFP_oriT*, and p*GFP* carry Tetracycline, Kanamycin, or Ampicillin resistance genes respectively.

**Supplementary Table 1.**
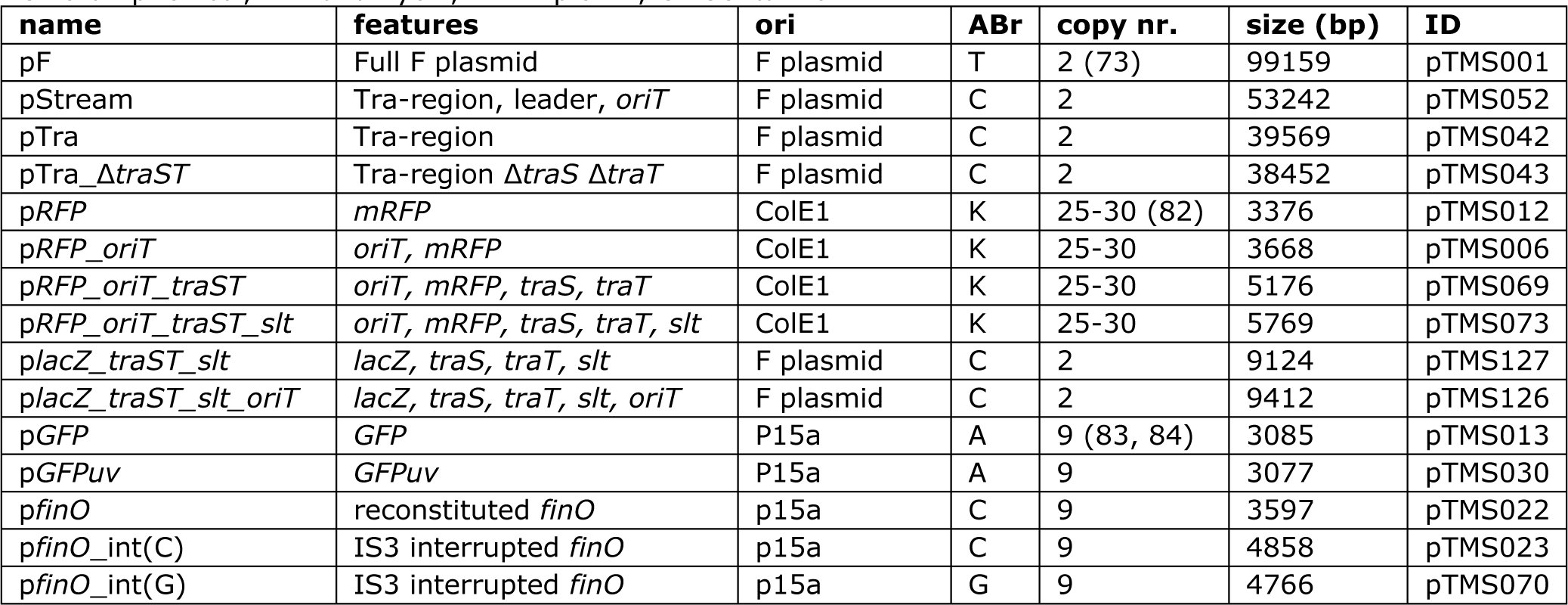
Plasmids used in this study. Plasmids used in this study, including the name as used in this chapter (name), genes and other DNA features they contain (features), their origin of replication (ori), antibiotic resistance (ABr), plasmid copy numbers (copy nr.), plasmid size (size (bp)), and the identifiers that have been used internally (ID). T: Tetracycline, C: Chloramphenicol, K: Kanamycin, A: Ampicillin, G: Gentamicin.

